# An early CYP26A1/CRYAA progenitor-glial domain marks the presumptive macular region during human retinal development

**DOI:** 10.64898/2026.07.17.738962

**Authors:** Qian Yang, Alberto Docampo-Seara, Michael B Powner, Gary Chung, Isabel Ribeiro Bravo, Elise Rody, Ryan B MacDonald, Sarah Giles, Krysztof Polanski, Carlos Talavera-Lopez, Kevin Eade, Sarah A Teichmann, Marcus Fruttiger

## Abstract

The human macula, a high-acuity retinal region essential for central vision, emerges early in development, but the cellular basis of its regional specialisation remains incompletely resolved. Building on recent studies that identified CYP26A1-mediated retinoic acid (RA) suppression in the presumptive macula, we define a CRYAA-positive progenitor-glial compartment in the temporal human retina from post-conception week (pcw) 7 onward. Single-cell RNA sequencing, immunohistochemistry and spatial morphometry indicate that this compartment is linked to regional gliogenic maturation and later corresponds to a specialised macular Müller glial population, rather than simply reflecting uniform pan-retinal developmental timing. Spatial mapping shows that the CYP26A1-positive domain remains delimited as the surrounding retina expands. Adult tissue analysis shows that CYP26A1 expression is retained in primate macular Müller glia and in corresponding acute-zone regions of visual-streak species. These findings extend current models of CYP26A1-mediated RA modulation in human acute zone/macular development by linking the early CYP26A1 domain to a regionally specialised CRYAA-positive progenitor-glial compartment. We propose that early macular patterning includes a spatially restricted glial programme that may contribute to later regional architecture and disease vulnerability.

## Introduction

High-resolution vision in humans critically depends on a small, specialised retinal area, the macula, from which most axonal projections to the brain originate. It contains tightly packed photoreceptors, mostly cones responsible for daylight colour vision, and retinal ganglion cells (RGCs), which are stacked up to eight cells deep. In contrast, the peripheral retina is dominated by rod photoreceptors, which support low-resolution monochrome night vision, and contains only a single layer of RGCs.

Other vertebrates have similar regional retinal adaptations for high-resolution vision, termed acute zones. The shape, size and location of the acute zone vary between species according to their visual-ecological needs (Baden et al., 2020). RGC density has been mapped in many animals and is commonly used to infer visual acuity across the retina and to define acute zones (Collin, 2008; Collin and Pettigrew, 1989; Hebel, 1976). Based on this approach, many mammals, including cats, dogs, pigs and rabbits, have a horizontally aligned acute zone referred to as visual streak. Visual streaks are also found in diurnal rodents such as hamsters and gerbils (Huber et al., 2010), but not in nocturnal mice and rats.

The relationship between animal acute zones, such as visual streaks, and the human macula is not well defined. Even in primates, which have a similar retinal anatomy to humans, comparison is not straightforward because of differences in eye size, raising the question of how the acute zone should be defined in primates and humans. The term macula is used only in humans and is clinically defined as a 5.5 mm diameter region. This area roughly matches the region where RGCs are multilayered but does not directly correlate with any other specific anatomical landmark.

Common blinding diseases such as age-related macular degeneration (AMD) and diabetic macular oedema predominantly affect the macula. However, the size of the diseased region can vary and may extend beyond the macula, depending on the patient and disease stage. In contrast, the less common macular disease macular telangiectasia type 2 (MacTel) clinically affects a defined oval region (around 3 x 2.5 mm) within the macula that is similar in size and shape between patients (Heeren et al., 2020), suggesting that this ‘MacTel area’ is anatomically or functionally distinct from the rest of the retina.

Regional differences in retinal disease susceptibility are likely to reflect regional differences in cell composition and support-cell specialisation that arise during acute-zone morphogenesis. Earlier work from our group reported early expression of candidate macular markers including CRYAA and CD44, followed by CRALBP expression during Müller glial differentiation in the temporal embryonic retina (Powner, 2011). Recent human and primate studies have since implicated CYP26A1-mediated modulation of RA signalling in early macular development and have shown that several candidate ‘foveal’ markers become enriched in macular Müller glia rather than defining a persistent foveal progenitor population (Harding et al., 2025; Hoshino et al., 2017; Krueger et al., 2024; Lu et al., 2020; Wohlschlegel et al., 2026; Zuo et al., 2024). These studies establish CYP26A1/RA modulation as a central feature of primate macular development. They do not fully resolve the cellular basis of the early CYP26A1-positive domain, its relationship to CRYAA expression and glial differentiation, or how this regional programme is retained as the eye expands. Here we examine these questions in human embryonic retina and adult mammalian retina. We define a spatially restricted CYP26A1/CRYAA-expressing progenitor-glial domain in the temporal retina that is consistent with early regional specialisation of macular Müller glia and with persistence of a related acute zone Müller glial identity in adult retina.

## RESULTS

### Asymmetry in the embryonic human eye

With the acute zone (macula) located in the temporal retina, the human eye is profoundly asymmetric along the nasotemporal axis. To study the emergence of this asymmetry, we analysed nasotemporal sections from human embryonic eyes from post-conception week (pcw) 7 to 12. This revealed pronounced morphological asymmetry between the temporal and nasal retina between pcw 8 and 11 (Fig. 1A-D), with the temporal retina being larger and showing a thicker nerve fibre layer, consistent with previous descriptions of accelerated temporal maturation (Hendrickson, 2016).

**Fig. 1.**
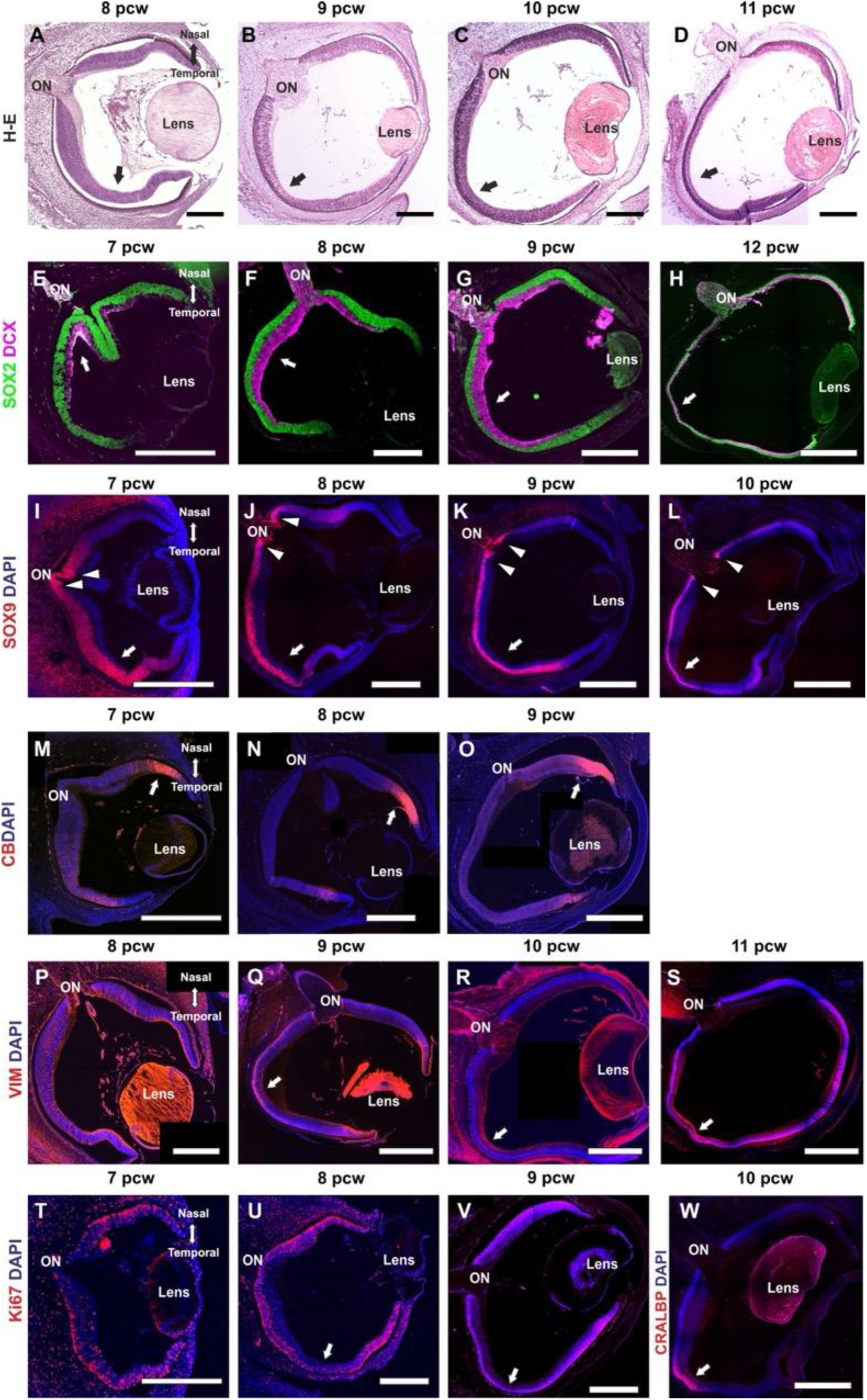
Early nasotemporal asymmetry in the human embryonic retina. (A-D) H&E-stained horizontal sections through human embryonic eyes at 8, 9, 10 and 11 pcw reveal pronounced asymmetry between temporal and nasal retina, with expansion of the temporal retina and a thicker inner retinal/nerve fibre compartment (arrows). (E-H) SOX2/DCX immunostaining at 7, 8, 9 and 12 pcw shows relatively symmetric SOX2-positive progenitor distribution, whereas DCX-positive differentiating inner retinal neurons and fibres are enriched in temporal retina (arrows). (I-L) SOX9 immunostaining at 7-10 pcw is stronger in temporal retina (arrows) and in retina adjacent to the optic nerve head (arrowheads). (M-O) CALB1 immunostaining at 7-9 pcw labels a peripheral retinal band posterior to the optic cup rim that is more prominent nasally (arrows). (P-S) Vimentin immunostaining at 8-11 pcw becomes stronger in temporal retina from pcw 9 onward (arrows). (T-V) KI67 immunostaining at 7-9 pcw shows relatively reduced proliferation in temporal retina (arrows). (W) CRALBP immunostaining at 10 pcw marks early differentiating Müller glia in temporal retina (arrow). Nasal-temporal orientation is indicated in the first panel of each relevant row. Scale bars: 250 µm (A, B, F, J, N, P, U), 500 µm (C, D, E, G, I, K, M, O, Q, T, V), 1 mm (L, R, W), 1.2 mm (S), 1.5 mm (H).

This raises the question of where the centre of the developing retina is and how this may be defined within an evolving retinal architecture. We used immunohistochemistry (IHC) with antibodies against SRY-box transcription factor 2 (SOX2) to visualise the outer retina and outer neuroblastic layer (ONBL), and doublecortin (DCX) to visualise the inner retina (Sánchez-Farías and Candal, 2015), consisting of the inner neuroblastic layer (INBL), the RGC layer and the nerve fibre layer (NFL) (Fig. 1E-H). SOX2 distribution, labelling retinal progenitors (RPs), showed no asymmetry in the developing retina and was not visible in the optic cup margin at pcw 7-9, but was strongly expressed in the ciliary marginal zone (CMZ) at the peripheral rim of the retina at pcw 12. In contrast, the inner retina, stained by DCX, showed pronounced asymmetry, with many more differentiating RGCs and nerve fibres in the temporal than in the nasal retina (arrows in Fig. 1E-H), a region previously considered the centre of the developing retina (Hendrickson, 2016).

SOX9 staining, which labels RPs and Müller cells (Poché et al., 2008), also showed an asymmetric distribution from pcw 7 onward, with more pronounced labelling in the temporal retina (arrows in Fig. 1I-L) and in retina adjacent to the optic nerve head (arrowheads in Fig. 1I-L). The centre of retinal SOX9 distribution is more temporal than the centre of DCX expression, demonstrating a difference between the inner and outer retina along the peripheral-to-central cell maturation axis. Another asymmetric pattern was seen with CALB1 staining, which labelled a band of peripheral retinal neuroepithelium posterior to the CMZ that was more pronounced in the nasal retina (arrow in Fig. 1M-O). Although CALB1 is typically considered a late neuronal differentiation marker, its robust expression here indicates an early, localised wave of post-mitotic maturation or a transient state associated with this early asymmetric development.

Vimentin staining, which marks RPs and glia, also showed increased intensity in the temporal retina containing the future macula/presumptive acute zone (PAZ) from pcw 9 onward (arrows in Fig. 1P-R). This was preceded by reduced proliferation from pcw 8 onward, as indicated by KI67 labelling (arrows in Fig. 1T-V), and is consistent with the previously described accelerated maturation of this region (Hoshino et al., 2017) and with CRALBP expression in differentiating Müller glia (Fig. 1W).

### Spatially differential transcriptional profiles

To further characterise asymmetry in human embryonic eye development, we micro dissected three retinal regions from both eyes from a pcw 8 and a pcw 11 embryo; 1) the peripheral rim, including the CMZ; 2) a more central nasal region (NAS); and 3) the temporal central retina, corresponding to the presumptive acute zone (PAZ). We analysed gene expression by bulk RNA-seq (Fig. 2A) from both eyes. Principal component analysis showed the strongest relationship between fellow eyes. CMZ samples (green in Fig. 2B) separated from the retinal samples and, within the retinal samples, grouping was affected more by embryonic age than by spatial position (Fig. 2B).

**Fig. 2.**
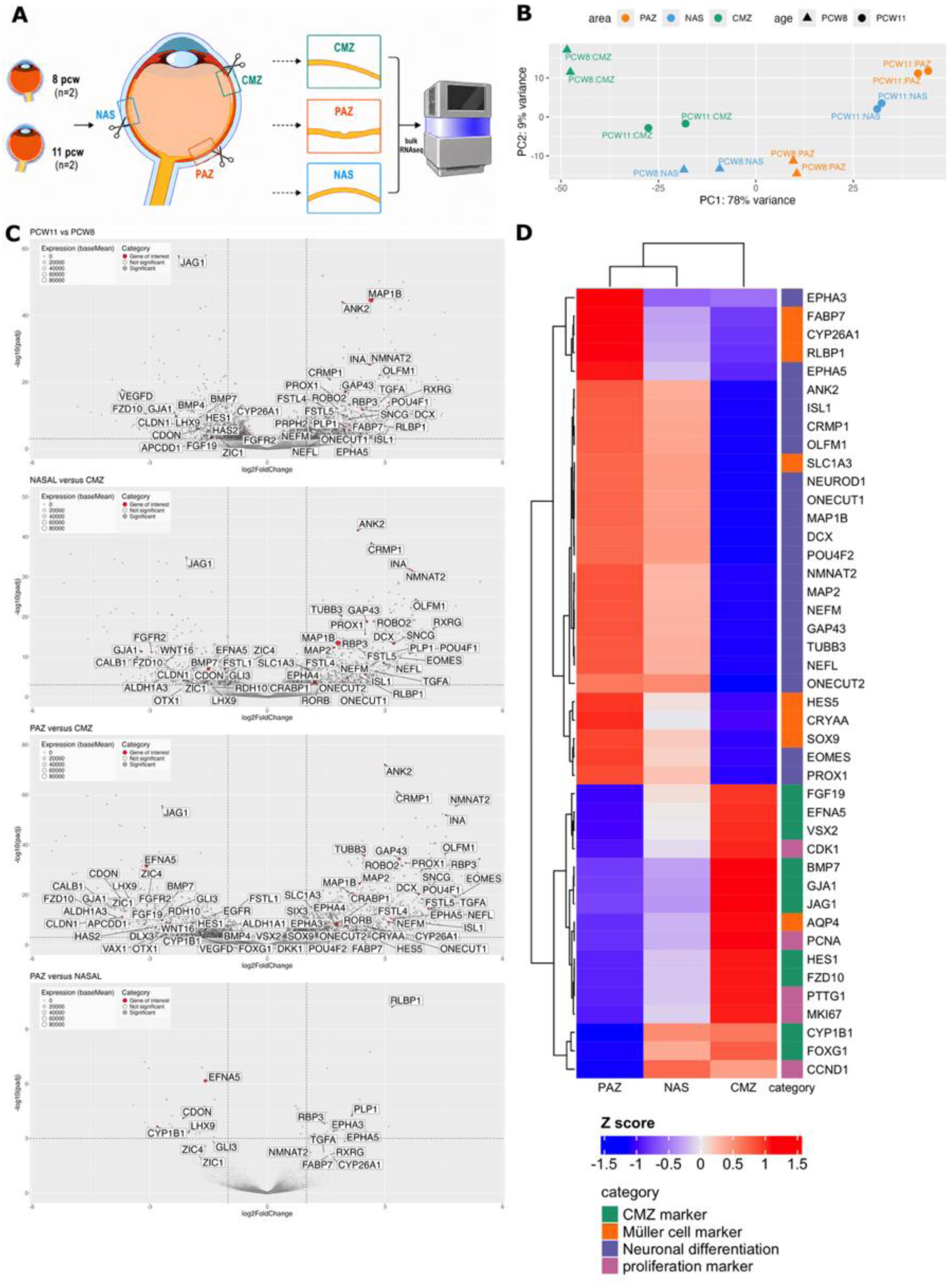
Bulk RNA sequencing reveals regional and developmental transcriptional differences in the human embryonic retina. (A) Schematic of the dissection strategy. Ciliary marginal zone (CMZ), nasal retina (NAS), and temporal presumptive acute zone (PAZ) were micro dissected from the eyes from pcw 8 and pcw 11 human embryos (2 embryos, 4 micro dissected eyes) for bulk RNA-seq. (B) Principal component analysis shows clear separation of CMZ from neural retina (PAZ and NAS), with developmental age (pcw 8 vs pcw 11) explaining a major source of variance. (C) Volcano plots of the indicated pairwise comparisons (pcw 11 versus pcw 8, NAS versus CMZ, PAZ versus CMZ, and PAZ versus NAS) highlight enrichment of neuronal differentiation genes in older and PAZ samples and enrichment of rim/CMZ markers in CMZ. Point size scales with expression level. Annotated significant marker genes are highlighted in red; significance was defined as fold change >= 2 and p < 0.001 for each pairwise comparison. (D) Heatmap of selected cross-region differentially expressed marker genes (CMZ, Müller glial, neuronal differentiation and proliferation) shows that PAZ exhibits a more advanced neuronal differentiation state than nasal retina and CMZ. Colour represents Z-scored average expression, averaged per region across pcw 8 and pcw 11 replicates.

Similarly, volcano plots of differentially expressed genes showed the strongest differences in comparisons between pcw 11 and pcw 8 samples, with increased expression of neuronal differentiation markers such as MAP1B and ANK2 and reduced expression of classical CMZ markers such as JAG1 and FZD10 in the older pcw 11 samples (Fig. 2C). A similar distribution of neural differentiation and peripheral rim markers was present in NAS versus CMZ and PAZ versus CMZ comparisons. Gene-expression differences between temporal and nasal retina were less pronounced but included increased neuronal differentiation markers, such as CRMP1 and NMNAT2, and decreased rim markers, such as JAG1 and CYP1B1, indicating greater maturity in temporal retina. Nasal markers (EFNA5 and FOXG1) and a temporal retinal marker (EPHA3) also showed clear differential expression, confirming correct dissection. In addition, Müller glial markers, including RLBP1 and FABP7, were enriched in temporal retina (Fig. 2C).

A heatmap of selected CMZ, neuronal and Müller glial markers further illustrates the advanced neural differentiation state of temporal retina, with neuronal markers (purple in Fig. 2D) more dominant in the PAZ-containing sample than in the nasal sample. The same markers are strongly reduced in the CMZ-containing sample. In contrast, CMZ progenitor markers (green in Fig. 2D) and proliferation markers (magenta in Fig. 2D) are enriched in the CMZ sample and reduced in the PAZ sample. Müller glial markers (orange in Fig. 2D) cluster with neuronal markers and are enriched in the PAZ samples.

To gain more detailed insight into gene expression and inferred lineage relationships at an early stage of eye development, we performed single-cell gene profiling with Smart-seq2 (SS2) on three micro dissected retinal regions (CMZ, NAS and PAZ) from both eyes from a pcw 8 (CS23) embryo. A total of 1,494 cells passed quality control and were included in the downstream analysis. Cells are visualised with force-directed graph embedding (FA) in Fig. 3A, showing 10 subpopulations with a main cluster of progenitor populations from which three differentiation trajectories emerge. In the central progenitor cluster, we identified three early-stage progenitor populations: one with neuroepithelial (NE) characteristics and two retinal progenitor populations (RPC0 and RPC1). The main differentiation trajectory emerging from this group was neuronal, with neuronal precursors (NP) and proliferating NPs feeding into a differentiating neuronal lineage that consisted mainly of RGCs, with a small number of photoreceptors (PRs) and horizontal cells (HCs). In addition, a Müller glial precursor (MGP) trajectory and a retinal pigment epithelium (RPE) precursor trajectory emerged from the central progenitor group (Fig. 3A).

**Fig. 3.**
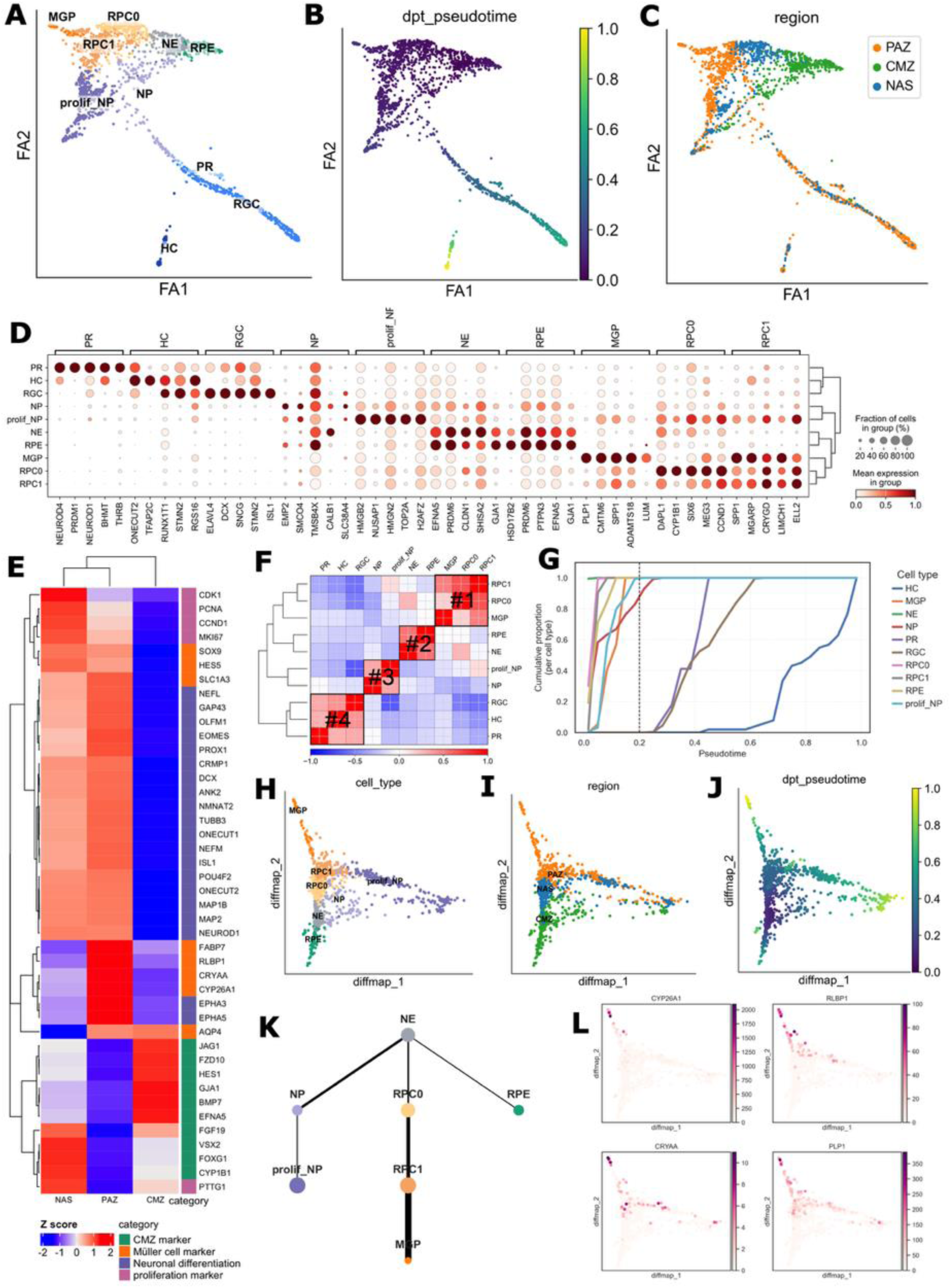
Single-cell RNA-seq identifies a PAZ-restricted gliogenic trajectory. (A-C) Force-directed graph embedding of 1,494 Smart-seq2-profiled cells from pcw 8 CMZ, nasal retina and PAZ. Cells are coloured by annotated cell type (A), pseudo time (B) and tissue of origin (C). The glial trajectory is detected only in the PAZ/temporal sample, whereas CMZ contributes the least differentiated progenitor-RPE populations. (D) Dot plot shows the average expression level and the fraction of cells expressing the top five markers for each cell group. (E) Heatmap of selected genes aggregated by dissection region from the single-cell dataset shows increased neuronal and Müller glial maturation in PAZ and progenitor enrichment in CMZ. (F) Cell-type correlation matrix identifies four major groups, including differentiating neuronal clusters and three progenitor-biased groups aligned with neurogenic, gliogenic and RPE-directed states. (G) Cumulative density plot showing how the proportion of each cell type changes with pseudo time; the vertical line at 0.2 marks where most of the progenitor cell wave ends. Progenitor cells from MGP, NE, NP, prolif_NP, RPC0 and RPC1 within pseudo time 0-0.2 were subset for analysis. Progenitor cells were viewed on diffusion map coloured by cell type (H), tissue of origin (I) and pseudo time (J). (K) PAGA trajectory showing the inferred lineage for NP, RPE and MGP. (L) Diffusion map showing selected markers for each lineage.

Pseudo time analysis showed that the neuronal differentiation trajectories were more mature than the larger cluster of less mature progenitors, proliferating neuronal precursors, MGPs and RPE precursors (Fig. 3B). Mapping the three dissection regions onto the FA plot confirmed that the neuronal trajectory was physically located in the nasal and temporal regions, with temporal cells (orange in Fig. 3C) more dominant among the most differentiated RGCs. This nasotemporal asymmetry was more pronounced in the emerging MGP trajectory, which was entirely derived from temporal retina. In contrast, the RPE trajectory was entirely derived from the CMZ-containing peripheral rim sample (green in Fig. 3C).

The five most significant marker genes for each cell population are shown in a gene-expression dot plot (Fig. 3D), showing population separation based on known cell type/state markers. In the more differentiated populations, these markers included well-established cell-type-specific genes such as NEUROD1 (PRs), THRB (cones), ONECUT2 (HCs), DCX and ISL1 (RGCs). The plot also included early RP and ciliary-margin components such as CCND1 and CYP1B1 (RPC0), ELL2 (RPC1), and markers of the ciliary margin-RPE interface such as CLDN1 and GJA1 (RPE). To further illustrate the distribution of established cell markers in this dataset, pseudo bulk expression was calculated for the three sample regions and plotted in a heatmap (Fig. 3E), using the same marker-gene selection as in Fig. 2D. This showed the same trends as the bulk RNA-seq data shown in Fig. 2. Progenitor and CMZ markers (green) were increased in the CMZ sample, and neuronal (purple) and Müller glial (orange) markers were enriched in the PAZ sample relative to both CMZ and NAS samples. Diffusion maps of selected marker genes for neuronal differentiation, progenitor status, proliferation and other markers further illustrate the link between the different subpopulations and the different regions (Suppl. Fig. 1).

Plotting the 10 cell populations in a correlation matrix (Fig. 3F) revealed four main groups: differentiating neurons containing RGCs, HCs and PRs (#4); committed neuronal precursors (#3); RPE and NE cells (#2); and RPs with MGPs (#1). Plotting the different cell populations over pseudo time (Fig. 3G) showed that the populations in group #4 (RGC, PR and HC) emerged substantially later than all other populations. To resolve the earlier populations more clearly, we used a pseudo time cut-off of 0.2 (Fig. 3G) and replotted the diffusion maps (using FA) with the remaining cells (Fig. 3H-J). In this reduced dataset, the NE population emerged more clearly as the least differentiated (Fig. 3J), and the three lineage trajectories (a neuronal path, an RPE path and a Müller gliogenic path) were more pronounced. A partition-based graph abstraction (PAGA) path plot also showed the three developmental trajectories emerging from the NE population (Fig. 3K). To further characterise the MGP lineage, we mapped expression of the PAZ marker CYP26A1 and the Müller cell marker RLBP1 in the reduced FA plots; both were enriched toward the distal tip of the MGP trajectory (Fig. 3L). Other genes that were notably enriched in the MGP lineage included PLP1 and CRYAA (Fig. 3L).

### Histological validation of lineage markers in the PAZ

To study the emergence of the MGP lineage histologically, we next used *in situ* hybridisation (ISH) and IHC to map the distribution of Müller cell markers beyond CRALBP (Fig. 1W). At pcw 10, transferrin (TF), one of the most highly expressed genes in adult Müller cells (Picard et al., 2008), was enriched in the PAZ compared with nasal retina. Glutathione peroxidase 3 (GPX3), which is expressed in adult, peripheral Müller cells (Velez et al., 2018; Yi et al., 2020), was enriched in the PAZ at pcw 16, indicating differentiating Müller cells in this region (Suppl. Fig. 2).

We also found PLP1 prominently expressed in the MGP trajectory, but PLP1 is not established as a marker for Müller cells or acute-zone development in the human retina. We therefore examined PLP1 distribution by ISH and found expression in the temporal ONBL at pcw 9, which spread into the nasal retina by pcw 10 (Suppl. Fig. 3). Of note, TF, GPX3 and PLP1 were also found around the optic nerve head in nascent retinal astrocytes, sometimes called peripapillary glia (Suppl. Fig. 2 and 3).

Another temporally enriched candidate Müller cell marker identified by RNA-seq was CRYAA. We assessed CRYAA by IHC and found that it was expressed at an earlier stage and in a more spatially restricted pattern than RLBP1, TF, GPX3 and PLP1 (Fig. 4A-I). Markers such as RLBP1 rapidly distributed pan-retinally after initial expression in temporal retina, whereas CRYAA remained restricted. In cross-sections, CRYAA appeared cytoplasmic and outlined entire cells with radial glia-like profiles. At pcw 8, CRYAA-positive cells had the simple, unbranched appearance of retinal progenitors (Fig. 4J), whereas older samples showed progressively more complex and branched morphologies (Fig. 4J-M), consistent with transition toward a Müller glial phenotype. Cells in the centre of the CRYAA-positive area showed a more differentiated appearance (Fig. 4L, L′) than cells at the edge of the CRYAA-positive area (Fig. 4L, L′′).

**Fig. 4.**
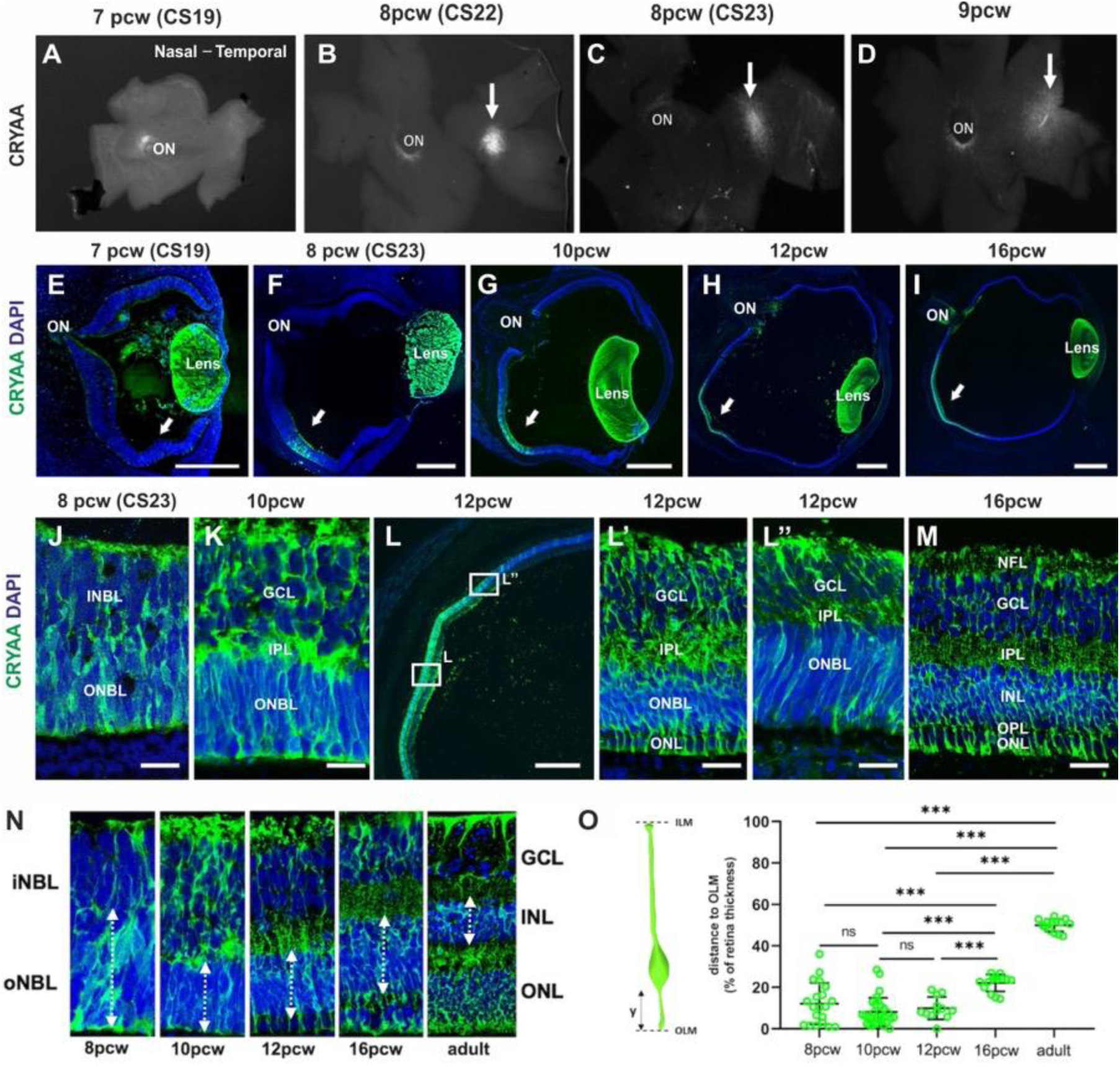
Localised CRYAA expression marks a sharply delimited temporal progenitor-glial domain. (A-D) Flat-mounted embryonic human retinae show a focal CRYAA-positive domain in temporal retina from pcw 7 to pcw 9 (arrows). (E-I) CRYAA immunostaining on sections at 7, 8, 10, 12 and 16 pcw reveals a spatially restricted temporal domain within the neural retina (arrows). (J-M) Higher-magnification views show age-dependent changes in CRYAA-positive cell morphology and laminar position: cells have simple radial progenitor-like morphology at pcw 8, acquire more elaborate radial profiles at pcw 10-12, and by pcw 16 occupy the inner nuclear layer with mature Müller glial morphology. Boxed regions in L are shown at higher magnification in L′ and L′′. (N) Representative CRYAA-positive cell profiles across development and in adult retina illustrate progressive inward displacement of cell bodies while radial processes are retained. (O) Schematic of the positional measurement (left) and quantification of CRYAA-positive soma position relative to retinal thickness (right) show little change from pcw 8-12 and a significant inward shift by pcw 16 and adult. Statistical annotations are *** p < 0.001. Scale bars: 1.5 mm (I), 500 µm (G, H, L), 250 µm (E, F), 50 µm (J, K, L′, L′′, M).

Because the temporal retina differentiates earlier than the rest of the retina, we asked whether CRYAA-positive cells were simply part of a regionally accelerated differentiation programme or instead represented an early regionalised progenitor-glial population. Morphometric analysis of the relative cell-body position of CRYAA-positive cells was consistent with an undifferentiated state between pcw 8 and pcw 12, with cell bodies localised within the outermost 40-20% of the retina. Only from pcw 16 onward did cell bodies show a statistically significant shift toward the inner retina, indicating Müller cell differentiation (Fig. 4N, O). This pattern suggests an early differentially regionalised progenitor state preceding Müller glial differentiation.

This concept was further supported by double labelling CRYAA-positive cells with the proliferation markers PCNA and Ki67 (Fig. 5A, Suppl. Fig. 5A-F). At pcw 8, the area of CRYAA expression overlapped with proliferation markers. Importantly, these cycling CRYAA-positive cells expressed SOX2 (Fig. 5B) and VSX2 (Fig. 5G), indicating that they were still retinal progenitors at this stage. This supports the interpretation that the CRYAA-positive spot represents a spatially restricted divergent domain in the temporal retina rather than simply the first step toward Müller glial differentiation in the PAZ.

**Fig. 5.**
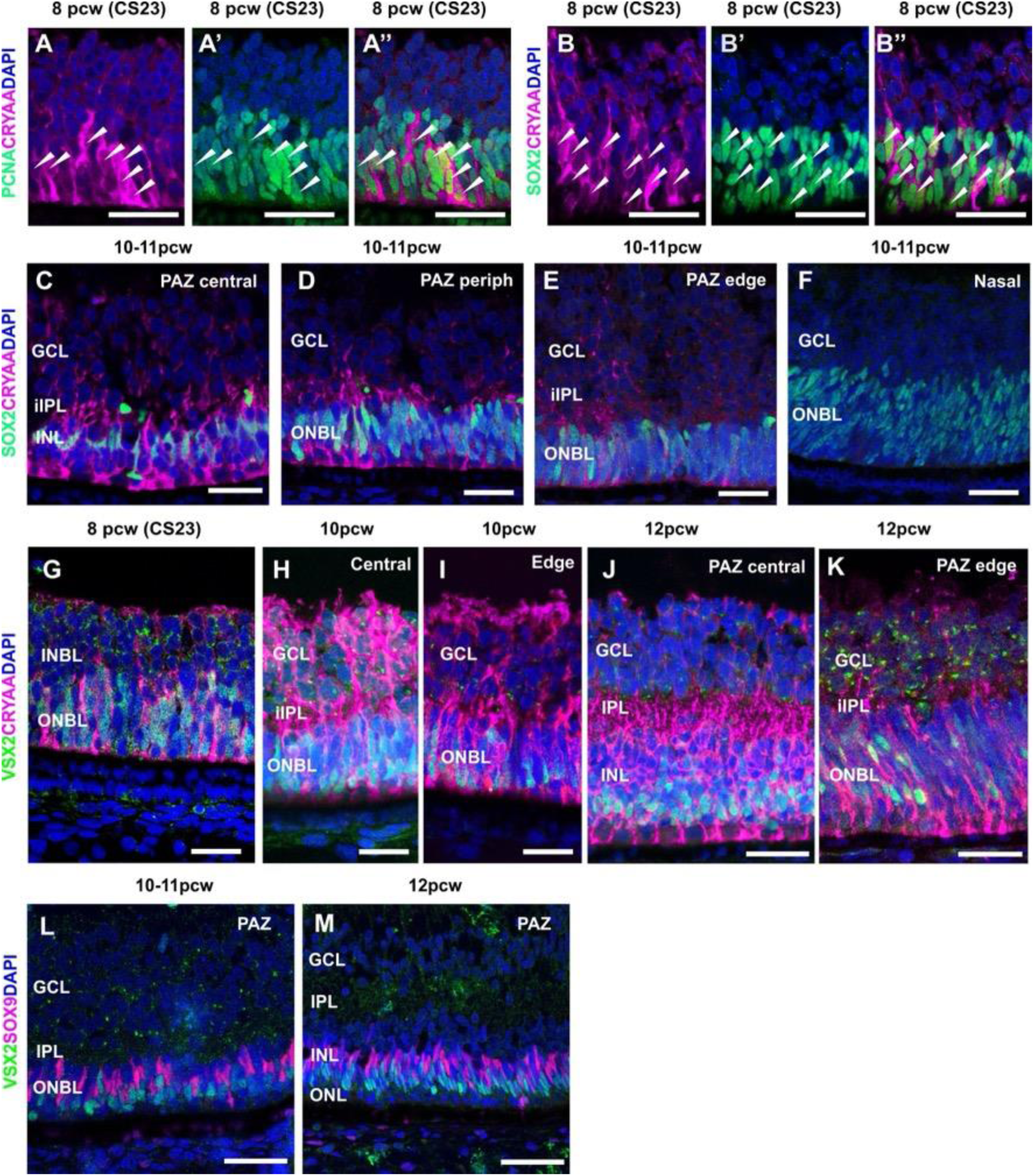
CRYAA-positive cells are cycling SOX2-positive progenitors that begin a central-to-peripheral gliogenic transition. (A-A′′) PCNA/CRYAA/DAPI staining in CS23 PAZ; A and A′ show single channels and A′′ the merge. Arrowheads mark CRYAA-positive/PCNA-positive cells. (B-B′′) SOX2/CRYAA/DAPI staining in CS23 PAZ; arrowheads mark CRYAA-positive/SOX2-positive cells. (C-F) SOX2/CRYAA/DAPI at 10-11 pcw comparing PAZ centre, PAZ periphery, PAZ edge and nasal retina. In the PAZ centre, some CRYAA-positive/SOX2-positive cell bodies are displaced slightly inward, whereas more peripheral PAZ and nasal cells remain in the outermost retina. (G-K) VSX2/CRYAA/DAPI at CS23, 10 pcw and 12 pcw show early colocalisation in outer retinal progenitors and, at later stages, a more mature central PAZ arrangement in which CRYAA-positive somata lie slightly inward of VSX2-positive nuclei; cells at the PAZ edge retain a more immature configuration. (L-M) VSX2/SOX9/DAPI at 10-11 and 12 pcw PAZ show SOX9-positive/VSX2-negative nuclei positioned inward of the VSX2-positive progenitor layer, consistent with emergence of a gliogenic trajectory. Scale bars: 50 µm.

Nevertheless, at pcw 10-11, the most central progenitors in the CRYAA-positive spot showed signs of differentiation, with subtle displacement of SOX2-positive cell bodies toward the inner retina, whereas more peripheral CRYAA-positive/SOX2-positive cell bodies remained in the outermost retina, similar to CRYAA-negative/SOX2-positive cells in nasal retina (Fig. 5C-F). From pcw 12 onward, CRYAA-positive cells in central temporal retina (PAZ) appeared quiescent, as also shown in Fig. 1T-V, and proliferation was detected only at the edges of the CRYAA- expressing retinal patch (Suppl. Fig. 5C-F). VSX2 staining at pcw 10 and 12 showed that VSX2- positive nuclei remained in the outermost position, whereas the CRYAA-positive soma lay further inward, consistent with Müller cell differentiation; more peripheral locations retained a more immature appearance (Fig. 5H-K). The emergence of a gliogenic trajectory in the ONBL at pcw 10-12 is indicated by SOX9-positive/VSX2-negative nuclei positioned inward of the VSX2- positive progenitor layer in the PAZ region (Fig. 5L, M).

Of note, we also detected CRYAA-expressing cells close to the optic nerve head from pcw 7 onward (Fig. 4A-D). These cells were adjacent to PAX2-positive nascent retinal astrocytes around the optic nerve head (Suppl. Fig. 4A-G). This discrete cell population occupied a SOX9-negative gap between SOX9-positive retinal progenitors and PAX2-positive/SOX9-positive astrocyte progenitor cells (Suppl. Fig. 4H-H′′) and was actively proliferating (Suppl. Fig. 4I-J). By pcw 16, the CRYAA-positive domain had expanded, but the small cuff of astrocyte progenitors around the optic nerve head remained negative. We also observed strong expression of TF, GPX3 and PLP1 in these cells (Suppl. Figs. 2 and 3).

### Retinoic acid signalling

Several studies have already identified CYP26A1 as an early marker of the future macula/fovea (Hoshino et al., 2017; Lu et al., 2020). Recent work has provided functional evidence that human macular/foveal development involves stage-specific CYP26A1-mediated modulation of retinoic acid (RA) signalling in the presumptive macula (Harding et al., 2025; Wohlschlegel et al., 2026).

In parallel, primate work has shown that CYP26A1 later localises to macular Müller glia, whereas FGF8 is not enriched in the foveal region (Krueger et al., 2024). Building on this framework, we used CYP26A1 as an established marker of the low-RA presumptive macular domain and asked how it is positioned within the expanding embryonic retina and how it relates to early glial differentiation. Spatial mapping of CYP26A1 expression over time (Fig. 6) showed a temporal spot from pcw 7 onward in an anterior position within the peripheral neural retina, proximal to the CMZ (Fig. 6A), in a region roughly defined by CALB1 expression at this early stage of retinal development (Fig. 1M). This region was limited to central sections containing the optic nerve.

**Fig. 6.**
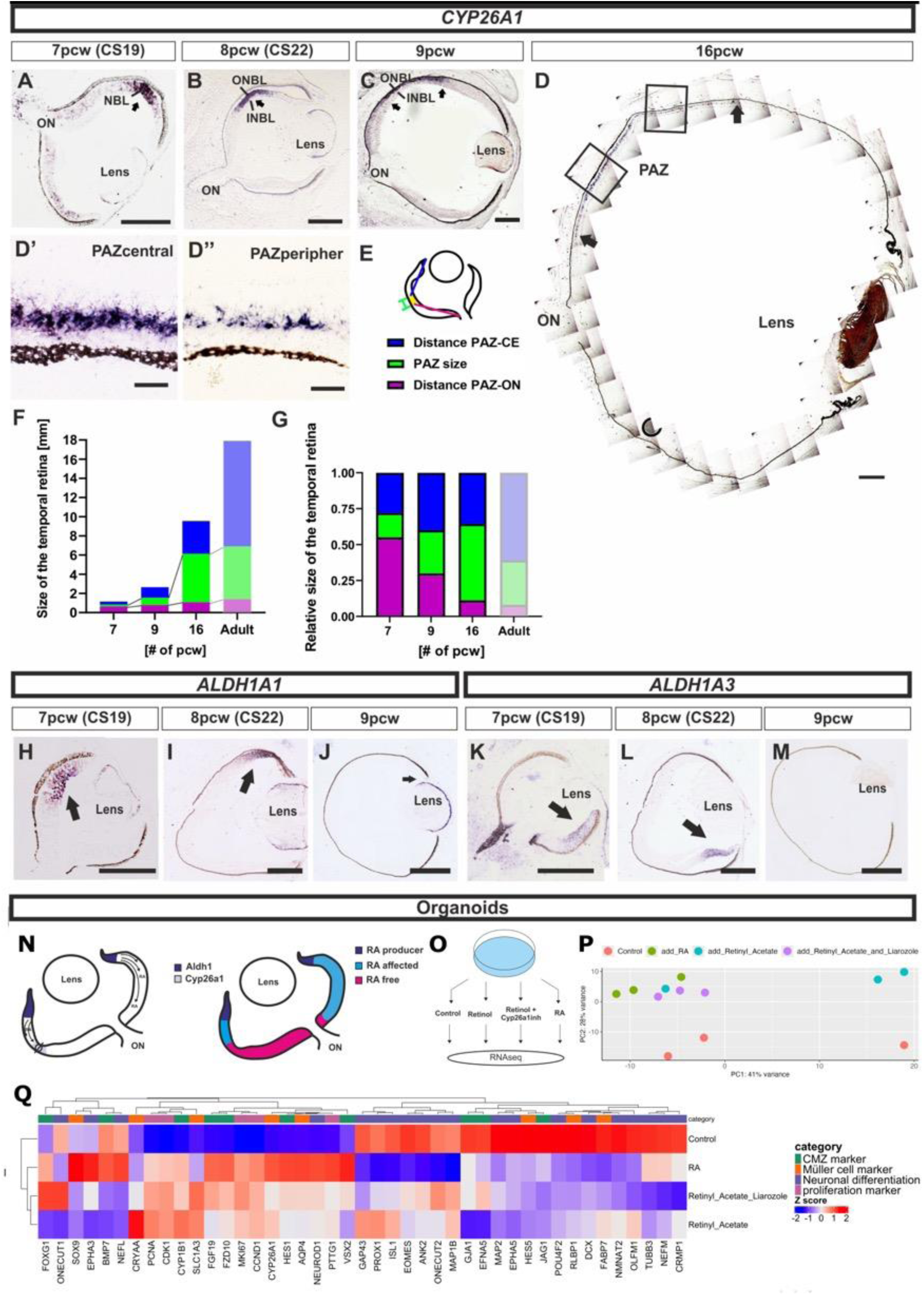
Spatial mapping of CYP26A1 and retinoic acid pathway components during development. (A-C) CYP26A1 *in situ* hybridisation on human embryonic eye sections at 7, 8 and 9 pcw shows a focal temporal CYP26A1-positive domain in anterior peripheral neural retina near the ciliary margin. (D) Reconstructed 16 pcw section showing the position of the presumptive acute zone (PAZ/CYP26A1-positive domain) within temporal retina. Boxed regions are shown at higher magnification in D′ (central PAZ) and D′′ (peripheral PAZ). (E) Schematic of morphometric measurements: distance from PAZ to the ciliary edge, PAZ size, and distance from PAZ to the optic nerve. (F-G) Absolute (F) and relative (G) quantification of these measurements at 7, 9 and 16 pcw and in adult retina shows that the CYP26A1-positive domain reaches near-adult size early while surrounding peripheral retina expands disproporti*onately.* (H-J) ALDH1A1 in situ hybridisation at 7-9 pcw labels dorsal peripheral *retina.* (K-M) ALDH1A3 in situ hybridisation at 7-9 pcw labels ventral optic stalk/retina. (N) Schematic showing the hypothesis underlying the organoid experiment: RA is produced in the CMZ, where ALDH1 expression is high, creating an RA gradient from CMZ to peripheral retina to PAZ, where CYP26A1 expression creates a low-RA zone. (O) Schematic of the experimental design. Organoids were treated with retinyl acetate (1 µg/ml), retinyl acetate plus liarozole (0.1 µM), RA (1 µM), or vehicle control. For each condition, three biological replicates underwent bulk RNA sequencing. (P) Principal component analysis of organoid RNA-seq samples shows treatment-dependent clustering. (Q) Heatmap of selected retinal progenitor, proliferation, Müller glial and neuronal differentiation markers (using the marker set shown in Fig. 2/3) shows induction of canonical RA-response genes and relative suppression of neural differentiation programmes under RA-elevated conditions. Scale bars: 1 mm (D), 250 µm (A, B, C, H, I, J, K, L, M), 25 µm (D′, D′′).

The relative position of the CYP26A1-positive region moved posteriorly over time (Fig. 6B-C), consistent with predominant retinal growth at the distal edge.

Quantification of the dimensions and position of the CYP26A1-positive region showed that it expanded in absolute terms and reached approximately 90% of its adult size by pcw 16. In relative terms, it moved toward a more central position because the retinal tissue distal to it expanded around 3.2-fold, whereas the distance to the optic nerve increased only 1.4-fold (Fig. 6F-G), matching observations by Hoshino et al. These morphometric data indicate that the CYP26A1-positive domain is established early and remains spatially restricted while the surrounding retina expands disproportionately. This behaviour is consistent with an early specified regional compartment in the future macula.

Because CYP26A1 degrades RA, we also visualised the enzymes that produce RA from retinaldehyde in our human eye samples. This confirmed the well-established mouse expression pattern of ALDH1A1 in dorsal retina and ALDH1A3 in ventral retina and optic stalk during embryonic eye development (Fig. 6H-M). We observed ALDH1A1 in dorsal peripheral retina and ALDH1A3 in ventral optic stalk and retina at pcw 7 and 8, as predicted from mouse expression patterns. Together with the CYP26A1 expression pattern, these data predict RA signalling in peripheral retina at pcw 7-8 and comparatively low RA signalling in the temporal CYP26A1- positive domain (Fig. 6N), in line with recent human macular datasets (Harding et al., 2025; Zuo et al., 2024).

Retinoic acid is required for normal eye development (Dupé et al., 2003). To test whether increased RA signalling broadly opposes early retinal differentiation, we exposed 5-week-old retinal organoids to three conditions expected to elevate RA signalling: (1) direct RA supplementation (1 µM), (2) retinyl acetate (1 µg/ml), and (3) retinyl acetate plus liarozole (a CYP26 inhibitor, 0.1 µM) (Fig. 6O). Gene expression was assessed by RNA-seq and showed qualitatively similar responses under all three elevated RA conditions compared to the control (Fig. 6P).

Canonical RA-response genes, including CYP26A1 and BCO2, were strongly induced, whereas ALDH1A3 was reduced, confirming activation of the RA pathway (Fig. 6Q). A heatmap (using the same markers as we used for Fig. 2 and 3) shows across all three conditions, proliferation-associated markers (such as PCNA, CDK1 and MKI67) increased and neuronal differentiation markers (such as DCX, CRMP1 and NMNAT2) decreased in response to RA. Consistent with recent organoid and foetal-retina perturbation studies (Harding et al., 2025; Wohlschlegel et al., 2026), these data support the view that elevated RA opposes aspects of early central retinal differentiation. In this study, these findings provide contextual support for interpreting the CYP26A1/CRYAA domain as a low-RA compartment influencing retinal development.

### CYP26A1 expression in adult retina

The temporal location of the embryonic CYP26A1-positive domain aligns well with the temporal position of the acute zone/macula in adult humans, suggesting that it marks a presumptive macular region during human eye development. By pcw 16, the size of this region and its distance to the optic nerve are already close to adult proportions, whereas subsequent enlargement of the eye depends predominantly on growth of the retina peripheral to this region (Hoshino et al., 2017).

Anatomically, the adult acute zone is based on differential cell distribution, which is also reflected in distinct metabolic signatures and gene expression across the retina (Bonelli et al., 2023; Peng et al., 2019; Yan et al., 2020; Yi et al., 2020). Single-cell studies by Peng et al., Yan et al. and Yi et al. assigned adult human/primate CYP26A1 expression to Müller glia and showed enrichment in the macular/foveal region. We mapped CYP26A1 distribution in adult human retina (Fig. 7A) and found expression in the inner nuclear layer in a region centred on the fovea and extending nasally to about the midpoint between the fovea and the edge of the optic disc (arrows in Fig. 7B). In cynomolgus macaque retina, the CYP26A1-expressing area was proportionally larger and extended to the optic disc (Fig. 7C-E), matching the proportionally larger anatomical acute zone in macaque. There was no expression in peripheral retina (Fig. 7F). In gerbil and pig retina, both of which contain visual streaks (Garcá et al., 2005; Huber et al., 2010), expression was found in a band in the superior retina that correlated well with the position of the visual streak (Fig. 7G-J). Overall, combining these prior single-cell assignments with our spatial mapping indicates that CYP26A1 expression is retained in primate acute-zone Müller glia and is also present in corresponding visual-streak regions of non-primate mammals.

**Fig. 7.**
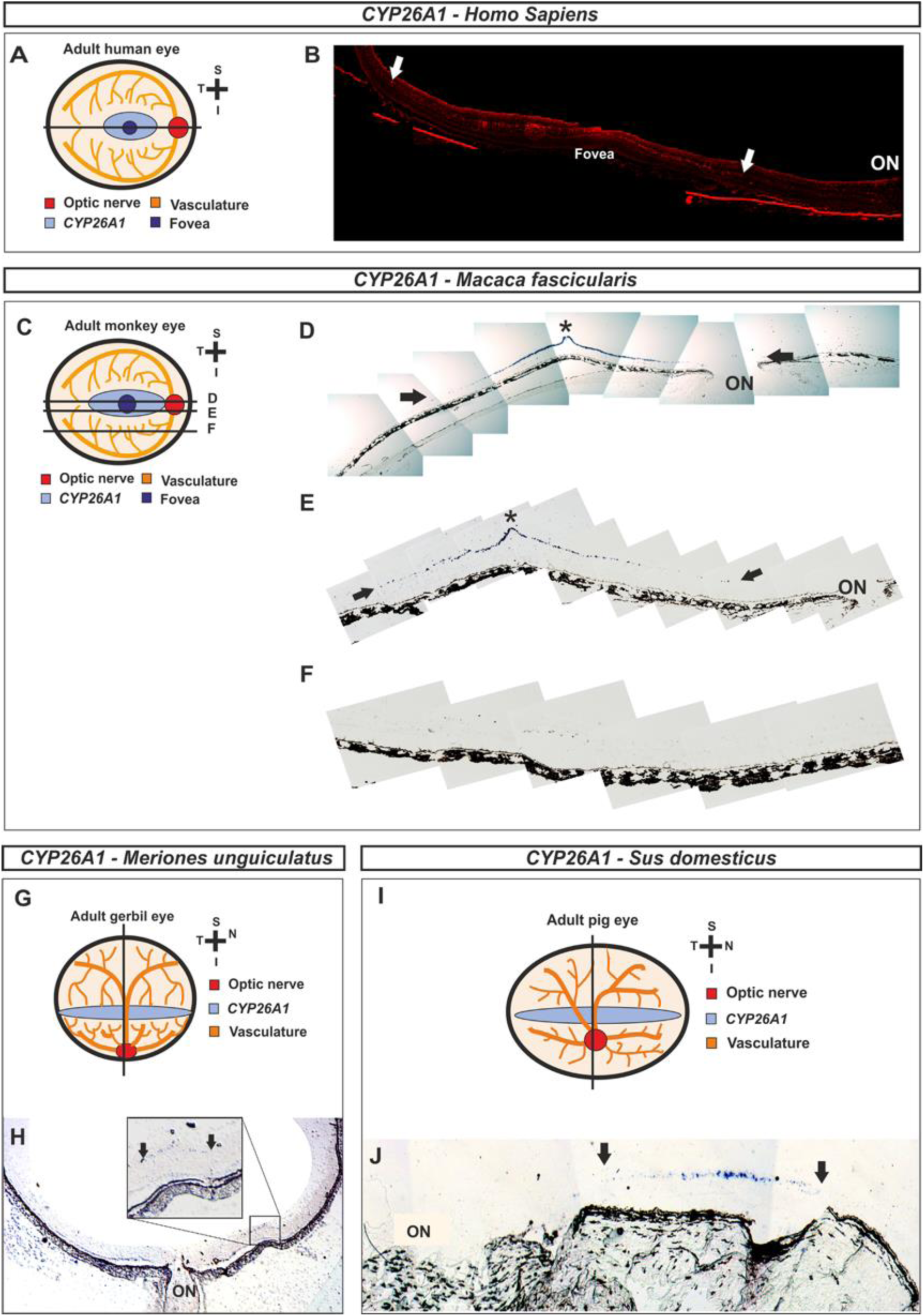
CYP26A1 expression identifies adult retinal acute-zone regions across mammals. (A) Schematic of the adult human eye showing the section plane. (B) Human retinal section showing *CYP26A1* labelling in the macular region, centred on the fovea and extending nasally toward the optic disc (arrows). (C) Schematic of the adult cynomolgus macaque eye showing section planes. (D-E) Stitched macaque retinal sections through the macular region show a broader *CYP26A1* - positive domain extending toward the optic nerve; asterisks indicate the foveal pit, and arrows mark the extent of the labelled region. (F) Peripheral macaque retina lacks detectable *CYP26A1* signal. (G) Schematic of the adult gerbil eye showing the section plane. (H) Gerbil retinal section demonstrates a band of *CYP26A1* labelling aligned with the visual streak (arrows; inset, higher magnification). (I) Schematic of the adult pig eye showing the section plane. (J) Pig retinal section demonstrates *CYP26A1* labelling in a superior streak-like band corresponding to the acute zone (arrows). Together with prior single-cell assignments of *CYP26A1* to primate Müller glia, these spatial data indicate retention of *CYP26A1* expression in primate acute-zone Müller cells and in corresponding visual-streak regions of non-primate mammals.

## DISCUSSION

In the present study we identify an early progenitor-glial cell population in the temporal human retina that aligns with development of the acute zone/macula. This domain is evident from pcw 7-8, expresses CYP26A1 and CRYAA, contains cycling SOX2-positive/VSX2-positive progenitor-like cells, and subsequently gives rise to a spatially restricted Müller glial population. The same CYP26A1-positive territory remains delimited as the eye expands and is retained in adult acute- zone Müller cells. These findings suggest that macular identity is not only a consequence of accelerated temporal retinal maturation or later neuronal packing, but includes an early, molecularly defined progenitor-glia programme.

Several recent studies have identified CYP26A1 and reduced retinoic acid (RA) signalling as features of the developing human or primate macula/fovea, and others have shown that several proposed foveal markers later map to macular Müller glia (Harding et al., 2025; Hoshino et al., 2017; Krueger et al., 2024; Lu et al., 2020; Wohlschlegel et al., 2026; Zuo et al., 2024). Our study builds on this work by defining the cellular substrate of the early CYP26A1-positive domain. The main distinction is that we link this territory to a CRYAA-expressing progenitor-glial compartment and to a Müller glial precursor trajectory, rather than treating CYP26A1 simply as a regional marker or RA readout. This places Müller glial regionalisation at the beginning of macular development.

The spatial design of the dataset was critical for this interpretation. By separately profiling micro dissected peripheral rim/CMZ, nasal retina and temporal presumptive acute-zone tissue, and then validating candidate markers histologically, we could distinguish general retinal maturation, nasotemporal developmental asymmetry, and macular/acute-zone regional identity. The temporal retina is developmentally advanced, but the CYP26A1/CRYAA domain is more restricted than a simple maturation wave. RLBP1, TF, GPX3 and PLP1 were enriched in temporal retina and are consistent with early Müller glial differentiation, but their expression patterns were broader and later than CYP26A1 and CRYAA. We therefore view these genes mainly as markers of a generic early Müller glial differentiation programme that appears first in temporal retina. CRYAA was different: it was present at the earliest stages in cycling SOX2- positive/VSX2-positive progenitor-like cells within the same retinal territory in which CYP26A1 was already expressed. Together, the spatial overlap, histology and trajectory data support an early CYP26A1/CRYAA progenitor-glial compartment without requiring direct co-expression in every cell of the domain.

CYP26A1 appears before clear molecular or morphological features of Müller gliogenesis, suggesting that a local RA sink is established before this domain differentiates. The simplest interpretation is that CYP26A1 contributes to the local conditions under which the compartment matures. This extends earlier work from our group identifying CRYAA, CD44 and CRALBP in the presumptive macula and proposing RA as a candidate morphogen (Powner, 2011), as well as human developmental transcriptomic studies that identified CYP26A1 as an early marker of the presumptive foveal/macular region (Hoshino et al., 2017; Lu et al., 2020), single-cell dual-omic data showing early regional specialisation (Zuo et al., 2024), and functional studies showing stage-specific RA suppression through CYP26A1 during human macula/fovea formation (Harding et al., 2025; Wohlschlegel et al., 2026).

Our organoid data offer supporting evidence, rather than proof, of CYP26A1/RA macular patterning. Conditions that increase RA signalling induced canonical RA-response genes and reduced neural differentiation markers, whereas the in vivo CYP26A1-positive domain corresponds to a region predicted to have lower RA tone. Together with published perturbation studies, these observations support a model in which reduced RA signalling favours local cell-cycle exit and early differentiation in the presumptive acute zone. RA signalling remains broadly required for normal eye development (Dupé et al., 2003), and its effects are stage-, dose- and tissue-context dependent. Our data therefore place the CYP26A1/CRYAA compartment within a low-RA developmental environment without implying that RA alone specifies macular identity.

A morphogenetic role of RA has previously been suggested in chick, where a high-acuity area is patterned by an RA-low/FGF8-high niche (da Silva and Cepko, 2017). In human and non-human primate retina, CYP26A1 appears to be the more robust feature, whereas FGF8 is not enriched in the presumptive macula/fovea and may remain mainly associated with the optic nerve head region (Harding et al., 2025; Krueger et al., 2024). Our findings fit the primate pattern. We observed CYP26A1 as a stable marker of the presumptive macular region, while the broader evidence argues against a simple transfer of the chick FGF8-centred model to primates. Of note, mice, which are nocturnal and lack an acute zone, express CYP26A1 in a streak in the superior embryonic retina but downregulate expression after birth (Sakai et al., 2004). A conserved feature may be embryonic local RA suppression, whereas the upstream signals and downstream patterning pathways differ between species.

However, the upstream signalling that initiates CYP26A1 expression in the temporal human retina remains unresolved in any species. In humans, the CYP26A1 domain and the CRYAA- positive progenitor-glial domain are already present by pcw 8, so our transcriptomic data are likely too late to capture the first instructive event. Several mechanisms remain plausible.

CYP26A1 could be induced by extrinsic signals from periocular mesenchyme, retinal pigment epithelium, optic stalk, lens or surface ectoderm. It could also reflect earlier axial patterning events established before or during optic fissure closure. A further possibility is that RA itself contributes to the induction of CYP26A1, creating a local negative-feedback domain that subsequently maintains low RA tone. Distinguishing between these possibilities will require earlier human material, primate or organoid models, and loss-of-function approaches that perturb CYP26A1 at the relevant developmental stage.

One mechanistic implication of our data is that Müller glia may provide a spatial scaffold for acute zone development. Müller cells span the thickness of the retina and, once differentiated, are relatively fixed in their radial and tangential positions. A regionally specified Müller glial compartment could therefore preserve positional information as the surrounding retina expands and as later-born neurons are added. In this framework, the CYP26A1/CRYAA progenitor-glial compartment may preserve positional information and shape the local signalling environment. Its role would not have to be direct specification of all macular cell types. It could bias cell-cycle exit, gliogenesis, neuronal differentiation and later tissue architecture within a fixed retinal territory.

Our morphometric analysis supports this interpretation and is consistent with a recent related analysis (Harding et al., 2025). The CYP26A1-positive domain is established early, expands in absolute size and approaches adult dimensions by pcw 16. At the same time, the retina peripheral to this domain expands much more than the distance between the domain and the optic nerve, consistent with an early specified region that remains spatially constrained while the eye grows around it. This provides a developmental explanation for how the macular region can retain a consistent position despite substantial post-patterning retinal expansion.

Our adult data extend this concept beyond embryonic development, supported by other human and primate single-cell data (Peng et al., 2019; Yan et al., 2020; Yi et al., 2020). In our sections, CYP26A1 expression persisted in the human macular region and in the macaque acute zone; and in gerbil and pig in the superior visual streak. Together, these data support CYP26A1 as a molecular marker of the acute-zone compartment, with a Müller glial assignment in primates and a corresponding acute-zone distribution in visual-streak mammals. This is useful because the term macula is anatomical and clinical rather than molecular. It is conventionally defined as a 5.5 mm region in humans, yet many central retinal specialisations and disease phenotypes occupy a smaller and more sharply defined area. CYP26A1 expression may therefore provide a more biologically grounded definition of the acute-zone compartment.

This molecular definition may be particularly relevant to macular telangiectasia type 2, which affects a stereotypical oval region that is similar in position and scale between patients (Heeren et al., 2020). The disease is also strongly linked to Müller cell pathology, with loss of Müller cell markers and Müller cell depletion (Powner et al., 2013, 2010). The overlap between the geometry of the MacTel area and the CYP26A1-positive acute-zone compartment raises the possibility that MacTel affects a developmentally specified macular Müller glial population, perhaps because acute-zone Müller glia retain distinct adult metabolic and signalling properties.

In summary, our findings support a model in which human acute-zone development involves early regionalisation at the progenitor level. A CYP26A1-positive temporal domain is established first and is followed by a CRYAA-positive progenitor-glial programme that gives rise to a specialised Müller glial compartment. Local CYP26A1 expression is predicted to lower RA signalling and may help promote early differentiation within this region. CYP26A1 expression is retained in primate acute-zone Müller glia and in visual streak regions of non-primate mammals. These findings argue that macular identity is not simply the consequence of accelerated temporal retinal maturation or neuronal packing, but includes an early, spatially restricted, molecularly defined progenitor programme that may shape normal acute zone architecture and regional disease susceptibility.

## Materials and Methods

### Tissue sample collection

Human embryonic and foetal material was provided by the Joint MRC/Wellcome Trust (grant# MR/R006237/1) Human Developmental Biology Resource (www.hdbr.org) and by the van Geest Centre for Brain Repair, Department of Clinical Neurosciences, University of Cambridge under ethics provided by HDBR (Project 200455). Adult human retina was provided by Moorfields Biobank at the UCL Institute of Ophthalmology Moorfields, approved by the Moorfields Biobank

Internal Ethics Committee and the NHS Research Ethics Committee (REC: 25/SW/0006) in accordance with the Human Tissue Act (HTA licence 12177). Informed written consent was obtained from all donors. Postmortem retinal tissue from adult macaque (Macaca fascicularis) was collected at the California National Primate Research Center (Animal Welfare Assurance ID: D16-00272 (A3433-01), Public Health Service Policy on Humane Care and Use of Laboratory Animals Policy). Postmortem eye tissue from Mongolian gerbil and domestic pig retinal tissues were obtained at UCL under institutional approval.

### Organoid cultures

Five-week-old retinal organoids were generated as previously described (Rosarda et al., 2023) and exposed for 2 days to the following factors (3 biological replicates were obtained for each condition): Retinyl acetate: 1ug/ml, retinoic acid (RA) 1µM, liarozole 0.1µM plus retinyl acetate 1ug/ml, and DMSO 1:1000 (control).

### Tissue processing for imaging

Human eye samples were used from embryos at CS19, CS22, CS23, and 9, 10, 11, 12, 16 pcw and adult. Furthermore, we also used postmortem eye tissue from adult macaque, gerbil and pig in this study. For ISH, samples were fixed and stored long term in PFA 4% at 4 °C until processed.

For IHC, samples were fixed in PFA 4% for 2h, then stored in PBS at 4 °C until processed. For cryosections, samples were cryoprotected in 30% sucrose and embedded in OCT compound (Tissue Tek, Torrance, CA) and froze at -80°C. Parallel series of sections were obtained in horizontal planes on a cryostat and mounted on Superfrost Plus slides (Menzel-Glasser, Madison, WI, USA). Samples were subsequently washed in Phosphate-Buffered Saline (PBS) and cryoprotected in 30% sucrose in PBS prior to embedding in OCT compound (Tissue-Tek) for cryo- sectioning. Serial sections (16 µm thick) were collected on Superfrost slides using a cryostat.

Sections were stored at -80 °C for *in situ* hybridization or at -20 °C for immunohistochemistry.

### Haematoxylin-eosin staining (H&E)

Haematoxylin-eosin staining was performed using standard protocols. Briefly, sections were washed in PBS to remove OCT and incubated with Mayer’s haematoxylin for 5-10 min. Excess haematoxylin was removed by rinsing the sections in running tap water for 10 min. Sections were then counterstained with eosin for 10 s and subsequently dehydrated through a graded ethanol series. Finally, samples were cleared in xylene and coverslipped using Eukitt quick-hardening mounting media (Sigma-Aldrich). Images were acquired using Leica DM IL inverted microscope (Leica, UK).

### Immunohistochemistry (IHC)

IHC was performed using standard protocols. Briefly, sections were washed in PBS at room temperature (RT) to remove OCT and incubated overnight at room temperature with the appropriate primary antibodies (SOX2, R&D systems AF2018, 1:500; DCX, Cell Signaling 4606, 1:300; Ki67, Dako GA626, 1:300; CB, 1:500; SOX9, Abcam ab185966, 1:500; Vim-Cy3, Sigma C9080, 1:300; CRYAA, Novus NB120-5595, 1:300; PAX2, Novus H00005076-M01, 1:300; PCNA, 1:500; CRALBP, ThermoFisher MA1-813, 1:300). Primary and secondary antibody incubations were performed in blocking buffer containing 4 % BSA and 0.5 % Tween in PBS.

For Ki67 and PCNA staining, antigen retrieval was performed prior to primary antibody incubation using a solution containing 90 % glycerol and 10 % sodium citrate (pH 6), heated to 120 °C and allowed to cool for 20 min at room temperature.

Following primary antibody incubation, sections were washed three times for 5 min in PBS and incubated with the appropriate Alexa Fluor-conjugated secondary antibodies (1:200) for 1 h at room temperature. Sections were then washed three times for 5 min in PBS and mounted with VectaShield HardSet mounting medium containing DAPI (Vector Laboratories, H-1500).

Images were acquired using an Invitrogen™ EVOS™ FL Auto 2 microscope (Thermo Fisher Scientific, UK) or a Zeiss LSM 700 confocal microscope (Zeiss, Oberkochen, Germany). Figures were assembled using CorelDRAW X8.

### *In situ* hybridization (ISH)

ISH was performed using the commercially available ViewRNA™ Tissue Assay (Thermo Fisher Scientific, UK) following manufacturer’s instructions.

Slides were pre-warmed at RT for 5-10 min and dehydrated sequentially in increasing concentration series (25 %, 50 %, 75 %, and 100 %) of ethanol for 10 min each, followed by dry baking at 60 °C for 1 h. Then, tissue digestion was performed using Proteinase K (provided in the ViewRNA™ kit, 1:100 dilution) for 5 min at 60 °C. Sections were then fixed in NBF (10 % formaldehyde in PBS) for 5 min at RT and rinsed twice in PBS before hybridization.

Hybridization was performed overnight at 40 °C with the corresponding probe of interest. Custom type 1 ViewRNA™ probes were obtained from Invitrogen and used at a 1:100 dilution. Slides were washed three times in washing buffer (ViewRNA™ kit) and incubated in PreAmplifier diluent (ViewRNA™ kit, 1:100) for 1 h at 40 °C. Slides were then washed three times for 2 min in washing buffer and incubated in Amplifier diluent (ViewRNA™ kit, 1:100) for 30 min at 40 °C. Samples were three times rinsed in PBS and incubated in Label Probe diluent (ViewRNA™ kit 1:500) for 1 h at 40 °C. After, slides were rinsed three times in washing buffer and incubated in Tris-NaCl buffer (pH 9) for 10 min at RT to proceed to signal detection.

Signal detection was developed using staining buffer containing 10% 1 M MgCl₂, 0.35% BCIP, 0.54% NBT, and 0.2% Tween in 10% PVA at 40 °C. Once optimal staining was achieved, slides were washed in Tris-NaCl buffer (pH 9.0) on a microplate shaker for 60 min and fixed in 4% PFA for at least 1 h. Finally, sections were dehydrated through a graded ethanol series and cleared in xylene shortly to be coverslipped with Eukitt quick-hardening mounting media (Sigma-Aldrich). Images were acquired using Leica DM IL inverted microscope (Leica, UK) or the bright field option of Invitrogen™ EVOS™ FL Auto 2 microscope (Thermo Fisher Scientific, UK).

### Deduplication and alignment of bulk RNA sequencing data

Raw FASTQ files were demultiplexed by firstly tagging the UMI (8-nucleotide sequence in R2 file) onto the R1 using UMI-tools (Version 1.1.5) on Galaxy Europe server (Version 1.1.5+galaxy1).

The output was then subject to quality control with FastQC (v0.11.9) and aligned to the human genome (gencode.v38) using STAR (v 2.7.11). The output bam file was deduplicated after sorting with Je-suite markdupes by setting using the default parameters and setting REMOVE_DUPLICATES="true". FeatureCounts (Subread v2.0.6) was then used for counting reads mapped to genes.

### Data processing and analysis of SS2 scRNA-seq data

Plate-based Smart-Seq2 (SS2) sequencing data were aligned against human reference GRCh38 using the STAR into featureCounts workflow followed by quality control steps: (1) cells with less than 300 genes detected were removed; (2) cells with less than 500 counts were removed; and (3) genes detected in fewer than 3 cells were removed. Preprocessing of the SS2 data was performed as following: normalization with pp.normalize_total (with 10,000 counts per cell after normalization), logarithmization with pp.log1p, highly variable genes detection with pp.highly_variable_genes, feature scaling with pp.scale (max_value=10), cell cycle analysis with tl.score_genes_cell_cycle, and regressed out technical variance with pp.regress_out(adata, [’total_counts’, ’pct_counts_mt’]), and principal component analysis with tl.pca(adata, svd_solver=’arpack’). Then further dimension reduction was performed with uniform manifold approximation and projection (UMAP) computed using tl.umap with 30 PCAs and 20 neighbours, with random_state=66. For clearer visualization of the lineage, we computed the force-directed graph embedding (FA) with sc.tl.draw_graph(adata, random_state=66, layout=”fa”). Python (v.3.13.3) and Scanpy (v. 1.10.4) were used.

### Clustering and annotation of SS2 scRNA-seq data

For unsupervised clustering, a neighborhood graph was constructed and Leiden clustering was performed using Scanpy (sc.tl.leiden, resolution = 1.2). Clusters exhibiting highly similar cell type-specific marker expression were subsequently merged. Cell identities were manually assigned based on the expression of established cell type-specific marker genes. Retinal ganglion cells (RGCs) were identified by expression of NEFM, NEFL, and GAP43; photoreceptors (PRs) by OTX2 and THRB; horizontal cells (HCs) by PROX1 and ONECUT2; Müller glial precursors (MGPs) by RLBP1; retinal pigment epithelium (RPE) by MITF, PMEL, and TYRP1; neuroepithelium (NE) by VSX2 and SOX2; and neuronal progenitors (NPs) by ASCL1. NP clusters exhibiting high S- and G2/M-phase cell cycle scores were annotated as proliferating neuronal progenitors (prolif_NP). Retinal progenitor cells (RPCs) were identified by expression of SIX6 and CCND1. Two transcriptionally distinct RPC populations, RPC0 and RPC1, were subsequently defined based on differential expression of key marker genes, including FGF19 and CYP1B1.

### Trajectory analysis of SS2 scRNA-seq progenitor subset

Pseudo time was first imputed in the entire dataset using sc.tl.dpt with neoroepithelial cells (NE) set as the origin. To zoom in on the retinal progenitors, only cells with pseudo time below 0.2 were included. Progenitor trajectories were inferred by combining PAGA and diffusion map on the progenitor subset. Pseudo time was then computed on this progenitor subset and reran diffusion map and PAGA with tl.diffmap and tl.paga, respectively. To visualize the development trajectories, diffusion maps were used with the position from PAGA with pl.embedding(adata_prog, "diffmap_").

## Acknowledgement

We thank Dr. Elisabeth Kugler for assistance with image analysis, Prof. Dr. Brian Clark for providing an initial analysis of the organoid dataset and Liv Remez for technical assistance. This work was supported by funding from the Lowy Medical Research Foundation and infrastructural support from the NIHR Moorfields Biomedical Research Centre.

---

**Supplemental Figure 1.**
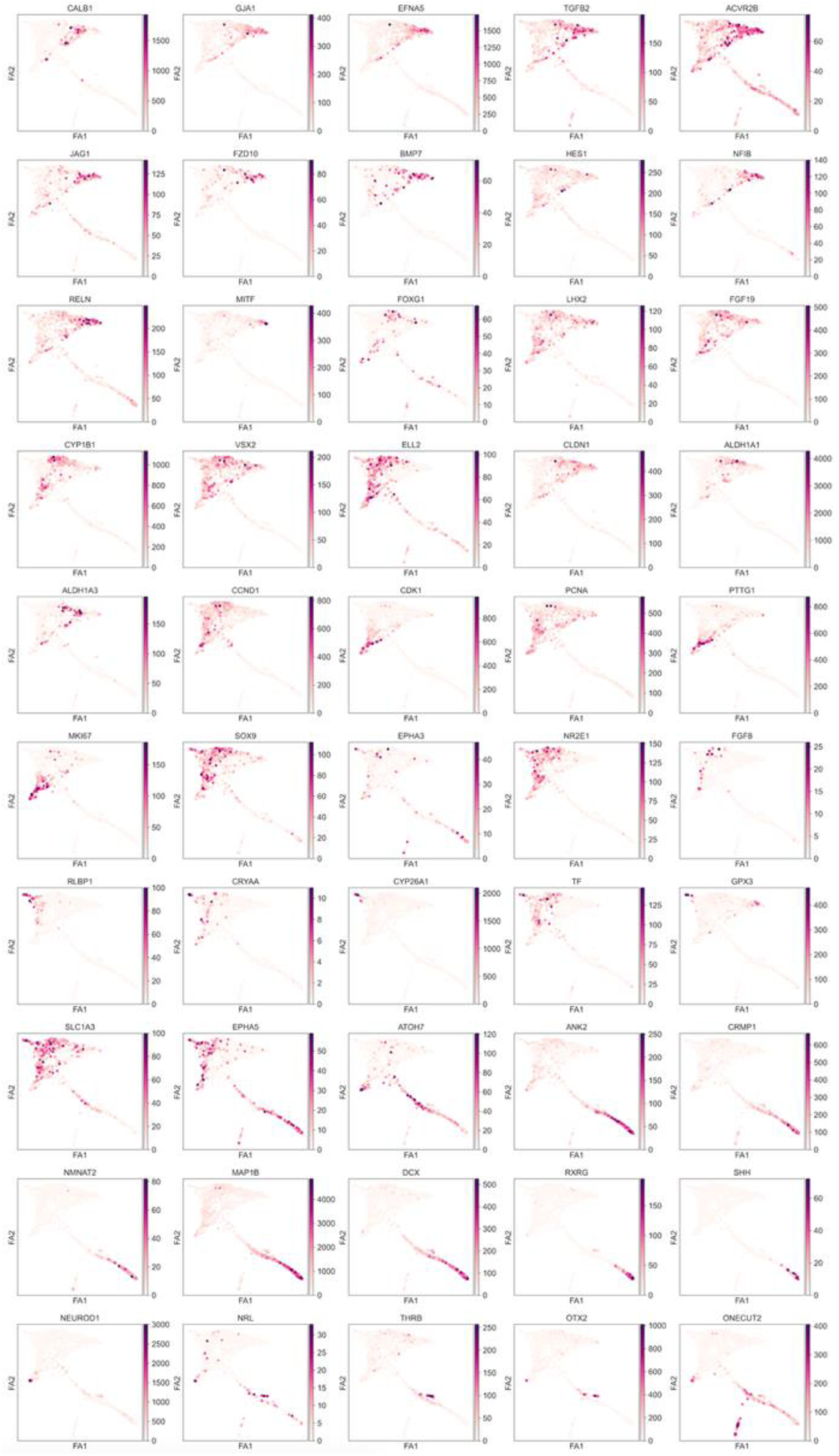
Expression of selected lineage, regional and cell-state markers across the pcw 8 retinal single-cell FA graph. FA graph embeddings of the 1,494 Smart-seq2-profiled cells shown in Fig. 3A-C, coloured by expression of the indicated genes. Markers include genes associated with early retinal progenitor and ciliary-margin/RPE states (including VSX2, CYP1B1, ELL2, CLDN1 and ALDH1A1/3), proliferation (CCND1, CDK1, PCNA, PTTG1 and MKI67), Müller gliogenic/PAZ identity (RLBP1, CRYAA, CYP26A1, TF and GPX3), and neuronal differentiation (ATOH7, ANK2, CRMP1, MAP1B, DCX, RXRG, NEUROD1, NRL, THRB, OTX2 and ONECUT2). Expression scales are shown separately for each gene; darker purple indicates higher expression. FA1 and FA2 denote the two force-directed graph coordinates.

**Supplementary Fig. 2.**
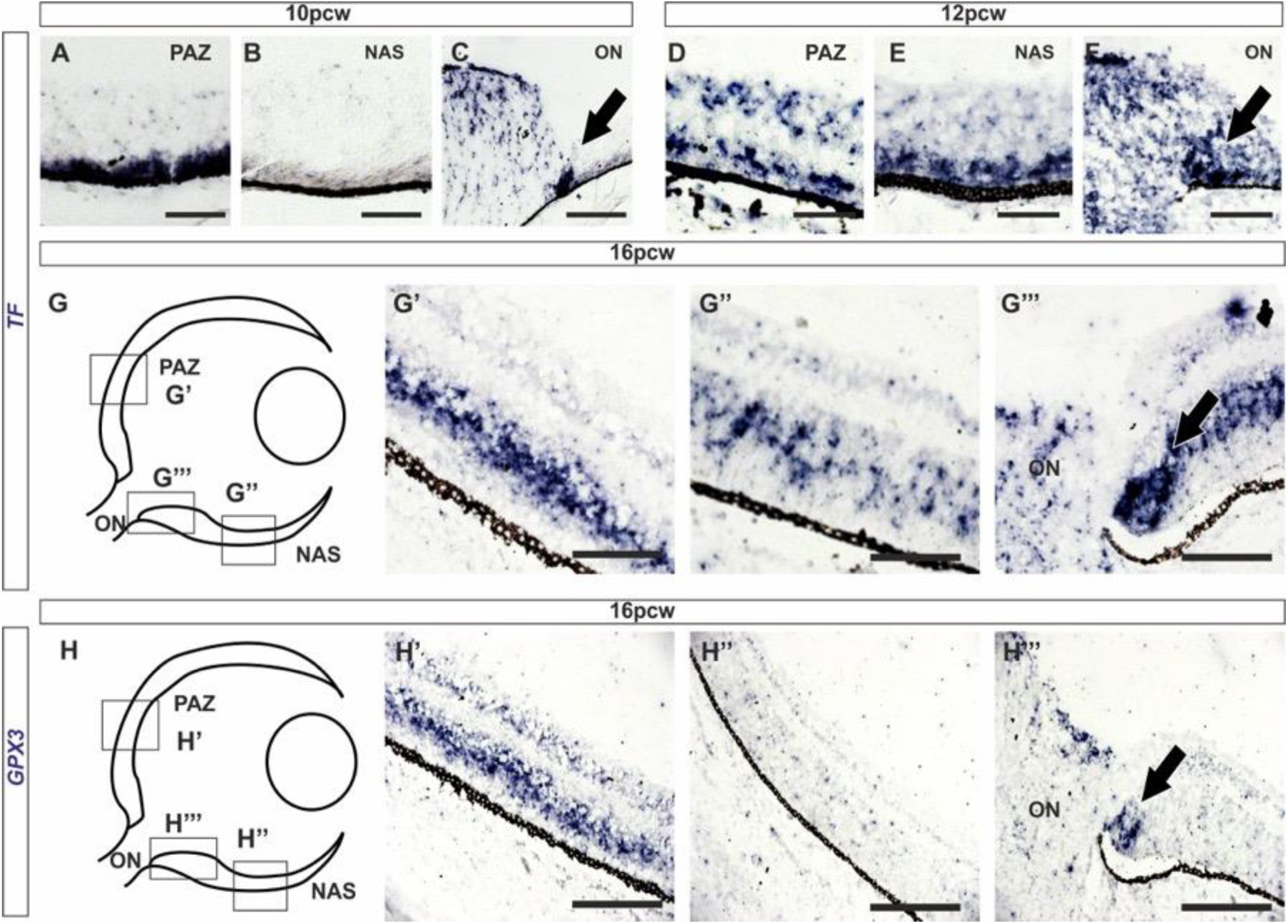
*TF* and *GPX3* are enriched in temporal retina and peripapillary glia during human retinal development. (A-C) *TF in situ* hybridisation at 10 pcw in presumptive acute zone (PAZ), nasal retina (NAS), and optic nerve head (ON) regions. (D-F) *TF in situ* hybridisation at 12 pcw in PAZ, NAS and ON. (G) Schematic of the 16 pcw section showing regions enlarged in G′-G′′′. At 16 pcw, *TF* is enriched in the PAZ/inner nuclear layer (G′), weaker in nasal retina (G′′), and also present around the optic nerve head in peripapillary glia (G′′′, arrow). (H) Schematic of the 16 pcw section showing regions enlarged in H′-H′′′. *GPX3* at 16 pcw is similarly enriched in the PAZ/inner nuclear layer (H′), reduced in nasal retina (H′′), and present around the optic nerve head/peripapillary glia (H′′′). Scale bars: 50 µm.

**Supplementary Fig. 3.**
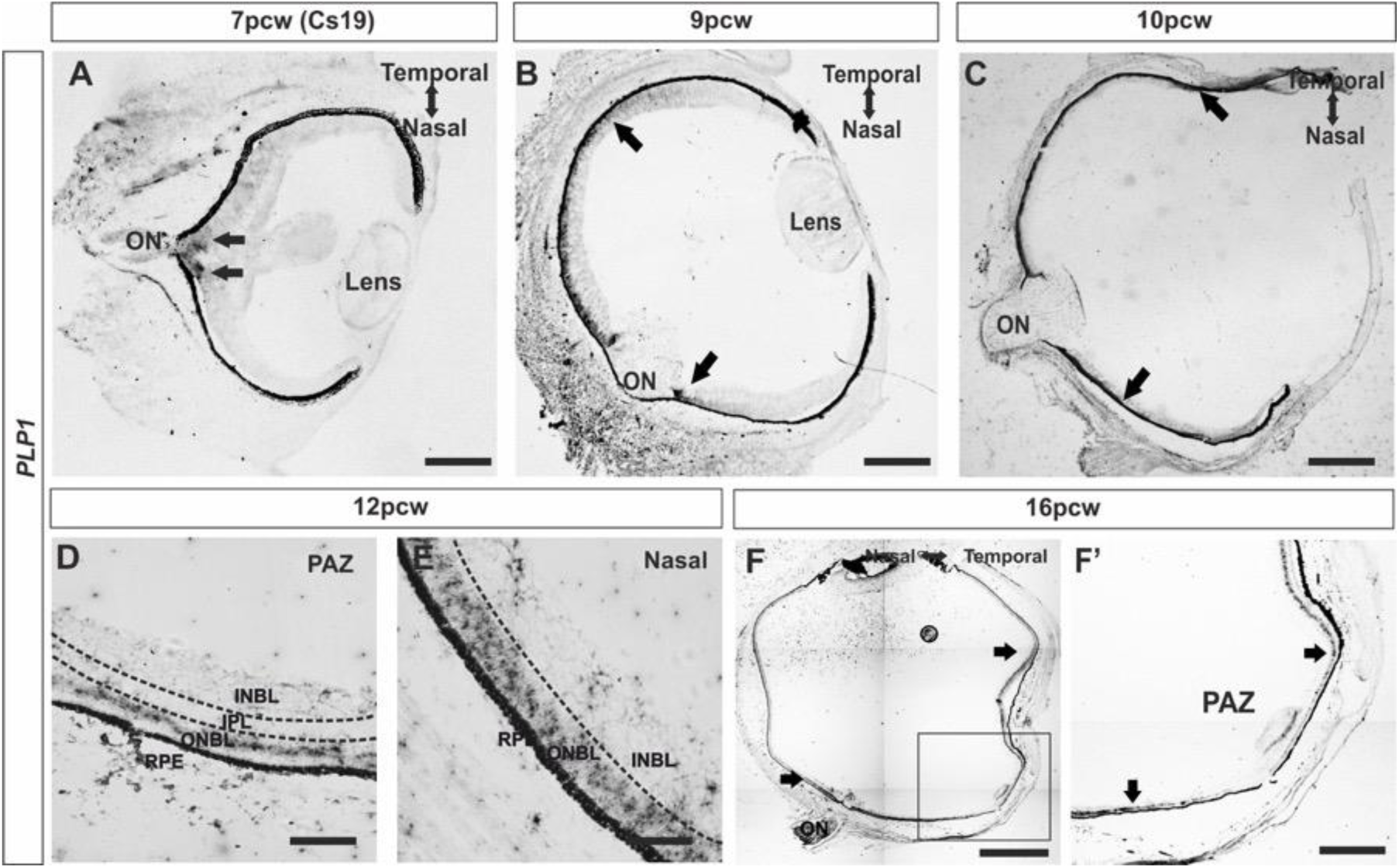
*PLP1* expression in temporal retina and peripapillary glia. (A-C) *PLP1 in situ* hybridisation on whole-eye sections at CS19/7 pcw, 9 pcw and 10 pcw shows signal in temporal retina and around the optic nerve head/peripapillary region (arrows). (D-E) Higher-magnification views at 12 pcw show stronger *PLP1* expression in PAZ than in nasal retina, with signal concentrated in the outer neuroblastic layer. (F) Whole-eye section at 16 pcw with boxed region enlarged in F′. By 16 pcw, *PLP1* is enriched in the PAZ and localised predominantly to the inner nuclear layer, consistent with progression toward a Müller glial state; additional PLP1-positive cells remain around the optic nerve head/peripapillary region. Scale bars: 1 mm (F), 500 µm (C, F′), 250 µm (A, B), 50 µm (D, E).

**Supplementary Fig. 4.**
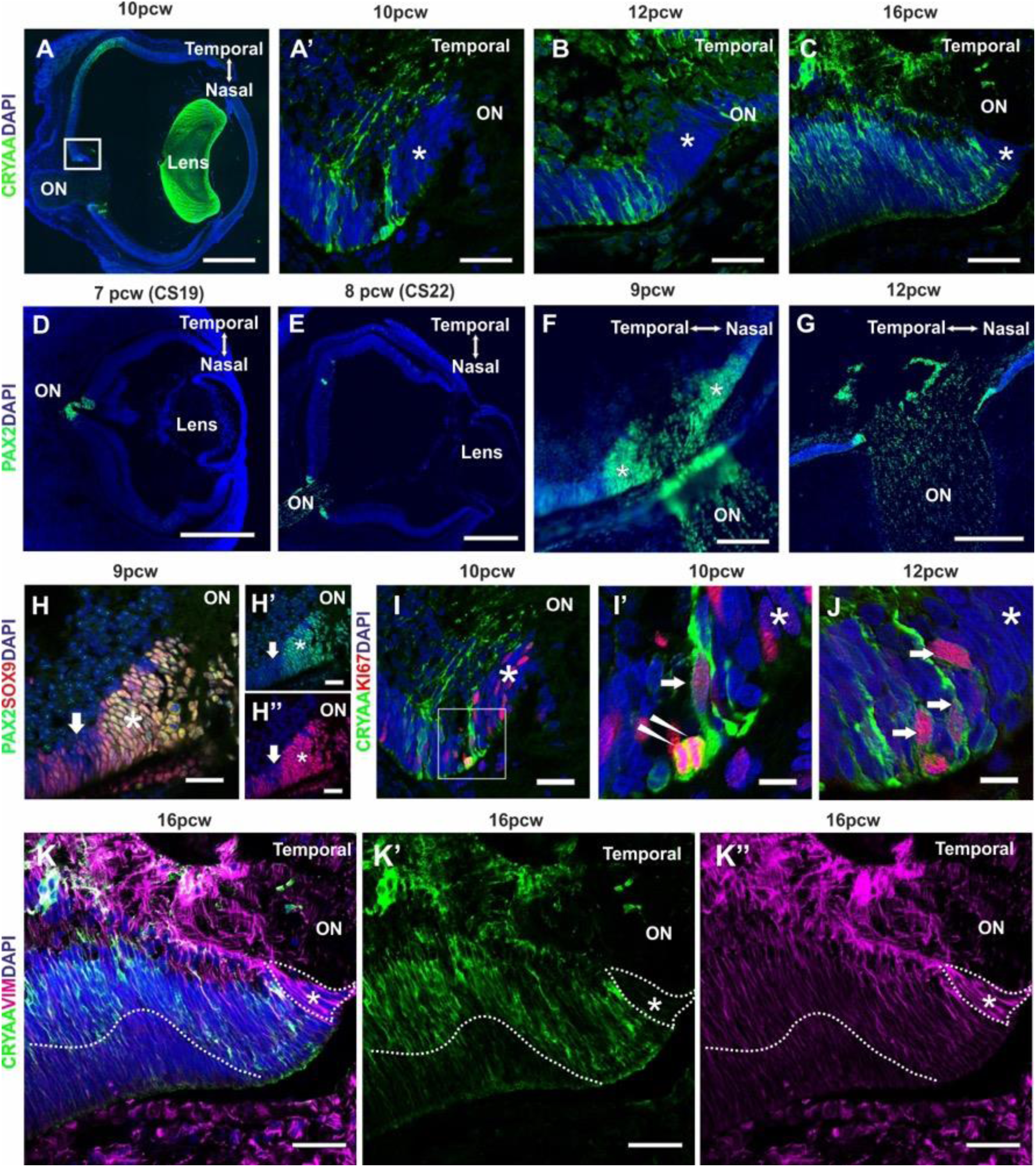
CRYAA is expressed around PAX2-positive nascent retinal astrocytes/peripapillary glia at the optic nerve head. (A) CRYAA/DAPI immunostaining at 10 pcw identifies CRYAA-positive cells at the optic nerve head; the boxed region is enlarged in A′. (A′-C) Higher-magnification views at 10, 12 and 16 pcw show CRYAA-positive cells/processes adjacent to the optic nerve. (D-G) PAX2/DAPI immunostaining at CS19, CS22, 9 pcw and 12 pcw shows nascent retinal astrocytes/peripapillary glia associated with the optic nerve head. (H-H′′) PAX2/SOX9/DAPI co-immunostaining at 9 pcw identifies a gap adjacent to the optic nerve (arrowheads) that is coincident with the CRYAA-positive cells. (I-J) CRYAA/KI67/DAPI immunostaining at 10 and 12 pcw shows that a subset of CRYAA-positive peripapillary cells remains proliferative; I′ is a higher-magnification view of the boxed region in I. (K-K′′) CRYAA/Vimentin/DAPI immunostaining at 10, 12 and 16 pcw shows CRYAA-positive cells embedded within the vimentin-positive glial scaffold surrounding the optic nerve head, with a gradient of differentiation toward the temporal retina. ON, optic nerve; *, astrocyte niche. Scale bars: 500 µm (A, D, E), 250 µm (F, G), 50 µm (A′, B, C, H, H′, H′′, I, K, K′, K′′), 15 µm (I′, J).

**Supplementary Fig. 5.**
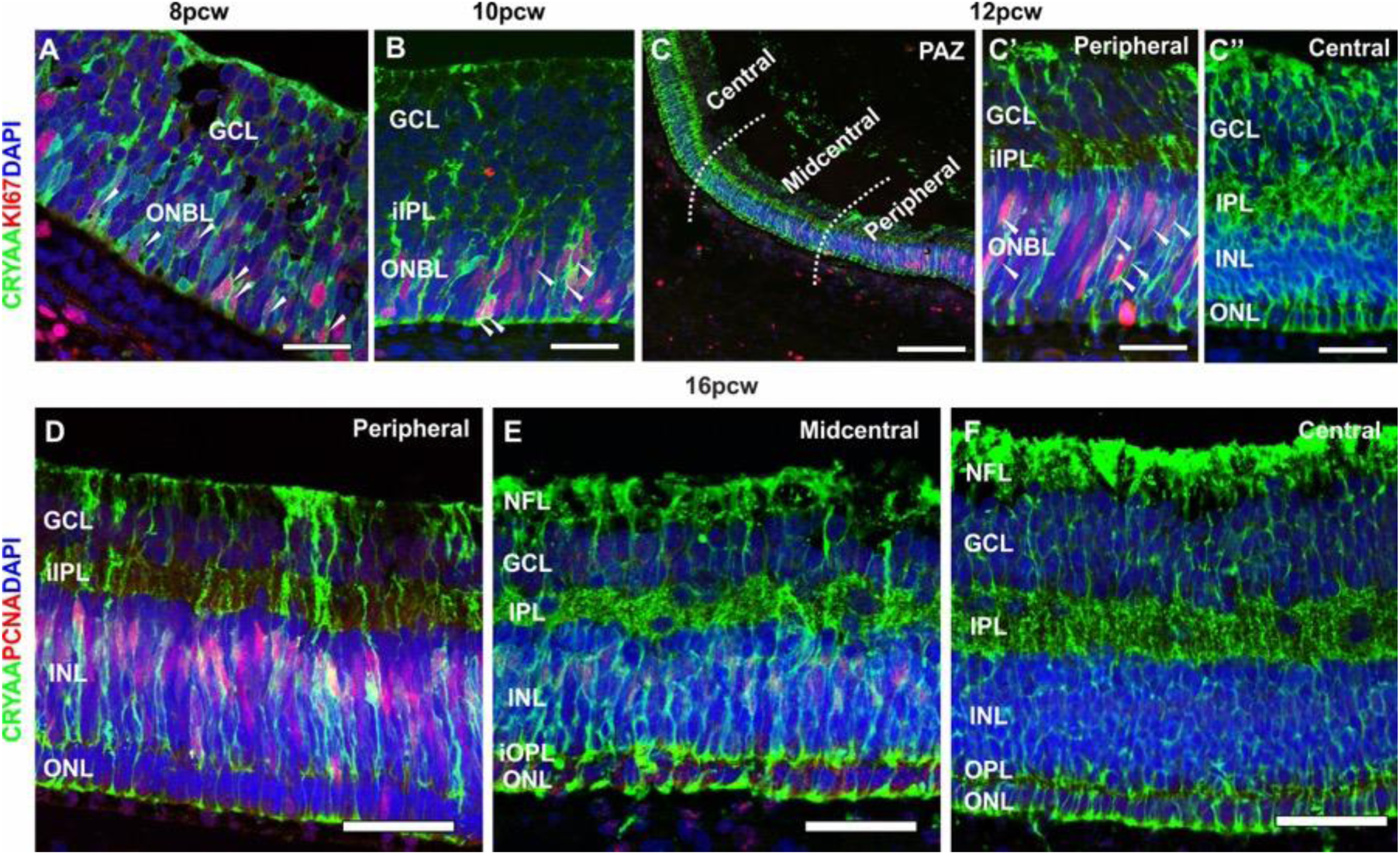
Proliferation of CRYAA-positive cells at later stages recapitulates early proliferation patterns in a gradient manner. (A-C′′) CRYAA/KI67/DAPI IHC from pcw 8 to pcw 12 shows that early proliferation patterns in CRYAA-positive cells (A, B) are present later in peripheral CRYAA-positive cells, whereas the most central cells are KI67-negative (C-C′′). (D-F) CRYAA/PCNA/DAPI IHC at pcw 16 shows a similar central-to-peripheral proliferation gradient to that observed at pcw 12 with KI67. Scale bars: 250 µm (C), 50 µm (A, B, C′, C′′, D, E, F).

